# Ferrets not infected by SARS-CoV-2 in a high-exposure domestic setting

**DOI:** 10.1101/2020.08.21.254995

**Authors:** Kaitlin Sawatzki, Nichola Hill, Wendy Puryear, Alexa Foss, Jonathon Stone, Jonathan Runstadler

## Abstract

Ferrets (*Mustela putorius furo*) are mustelids of special relevance to laboratory studies of respiratory viruses and have been shown to be susceptible to SARS-CoV-2 infection and onward transmission. Here, we report the results of a natural experiment where 29 ferrets in one home had prolonged, direct contact and constant environmental exposure to two humans with symptomatic COVID-19. We observed no evidence of SARS-CoV-2 transmission from humans to ferrets based on RT-PCR and ELISA. To better understand this discrepancy in experimental and natural infection in ferrets, we compared SARS-CoV-2 sequences from natural and experimental mustelid infections and identified two surface glycoprotein (Spike) mutations associated with mustelids. While we found evidence that ACE2 provides a weak host barrier, one mutation only seen in ferrets is located in the novel S1/S2 cleavage site and is computationally predicted to decrease furin activity. These data support that host factors interacting with the novel S1/S2 cleavage site may be a barrier in ferret SARS-CoV-2 susceptibility and that domestic ferrets are at low risk of natural infection from currently circulating SARS-CoV-2. This may be overcome in laboratory settings using concentrated viral inoculum, but the effects of ferret host-adaptations require additional investigation.

## Introduction

Severe acute respiratory syndrome coronavirus 2 (SARS-CoV-2), the virus that causes COVID-19, is a zoonotic member of *Coronaviridae* that emerged in 2019 as a major viral pandemic (1). As of August 2020, there have been over 20 million confirmed COVID-19 cases globally and approximately 761,000 deaths (2). SARS-CoV-2 uses angiotensin I converting enzyme-2 (ACE2) as its primary cellular receptor for host entry and infection (3-5). *In silico* analyses of ACE2 genes in diverse mammalian species have shown that residues important to viral binding are moderately conserved between humans and several domestic animals, and a broad range of species have been demonstrated to be permissive to infection *in vitro* and *in vivo* (6-10).

It is not yet known if natural infection of animals plays a role in public health epidemiology or has the potential to establish endemic reservoirs and threaten wildlife. SARS-CoV-2 has been observed to be capable of natural human-to-animal reverse-zoonoses, transmitting from infected individuals into mink (11), dogs (12) and felines (13-15). American mink (*Neovison vison*) are currently the only species observed to have natural human-to-animal spillover and onward transmission (11). To date, at least 27 mink farms in the Netherlands, Spain, Denmark and United States have reported outbreaks, including at least one probable case of mink-to-human transmission (16, 17). SARS-CoV-2 has also been shown to productively infect several species including ferrets and domestic cats *in vivo* (9, 10, 18, 19). Ferrets (*Mustela putorius furo*) are of special relevance to laboratory studies of respiratory viruses like *Influenza A virus* and recapitulate clinical pathophysiological aspects of human disease. Given their susceptibility to experimental infection and onward transmission via direct and indirect contact, ferrets have been proposed as an animal model to study SARS-CoV-2 transmission. Based on *in vivo* data, we expect all naïve ferrets in direct contact with an infected ferret will 1) become infected and 2) have measurable viral shedding or RNA via oral swabs up to 19 days post-infection and 3) seroconvert with measurable antibodies against SARS-CoV-2 receptor binding domain (RBD) (18, 19).

In March 2020, during the first wave of the SARS-CoV-2/COVID-19 pandemic in the New England area, we developed a rapid response study to investigate the potential for human-to-animal spillover and onward transmission in domestic, farm and wildlife species (CoVERS: Coronavirus Epidemiological Response and Surveillance). The goal of CoVERS is to understand if and how SARS-CoV-2 transmission is occurring at these interfaces to refine public health guidelines, investigate if there are additional risks to animal or human health associated with spillover and evaluate the potential for establishment of endemic reservoirs. Here, we highlight one enrolled household that created an exceptional natural experiment with direct relevance to our understanding of SARS-CoV-2 reverse zoonosis and animal models of disease.

## Results

### Absence of natural SARS-CoV-2 human-to-ferret transmission in a high exposure setting

A household with 29 free-roaming ferrets cared for by two adults was enrolled in the CoVERS study. Individual 1 experienced fever and fatigue from March 25-April 6 and Individual 2 experienced a sore throat, anosmia, migraine and fatigue from March 28-April 13 (Fig. 1A). Individual 2 tested positive for SARS-CoV-2/COVID-19 infection by nasopharyngeal swab and RT-PCR on April 1. Individual 1 is a probable positive due to the timing and symptoms but was not tested. Neither person was hospitalized, and both cared for the ferrets during the entirety of their disease courses.

**Figure 1.**
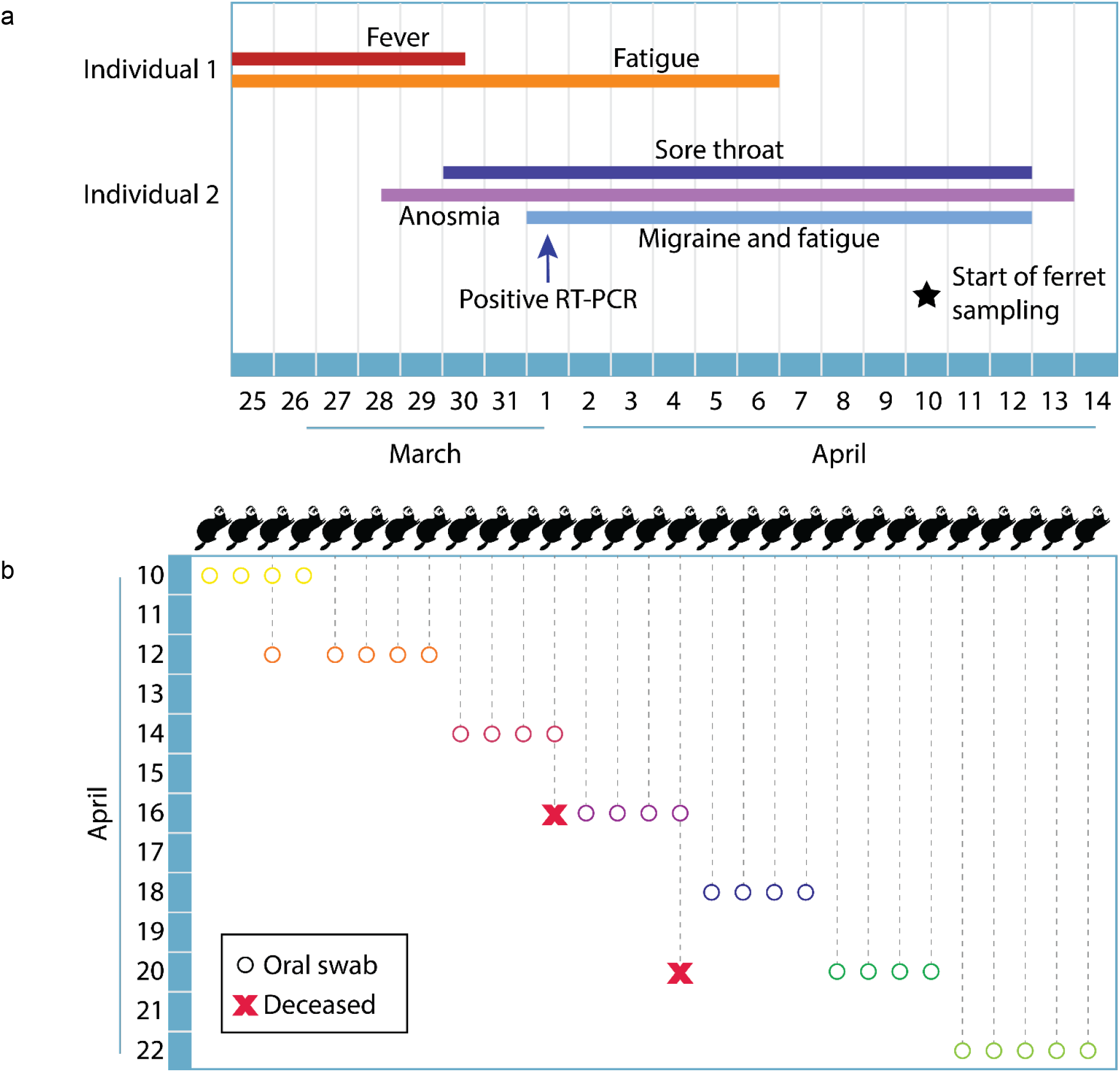
COVID-19 disease course and ferret sample collection timeline. A household with two adults and 29 free-roaming ferrets was enrolled in the CoVERS study. Both adults exhibited symptoms of SARS-CoV-2 infection in late March to early April of 2020, and one tested positive by RT-PCR on April 1^st^ (a). Oral swabs were collected from all ferrets in the home over a two-week period, beginning April 10^th^, concurrent with symptomatic disease in Individual 2 (b). One ferret (3) was sampled twice. Two 7-year-old ferrets (12 and 16) died during the study period, one by euthanasia due to chronic disease, the other cause is unknown.

A two-week, in-home sample collection scheme was designed to begin during the household quarantine period (Fig. 1B). The ferrets were free to move in all spaces of the home during this period and handled as usual, including regular petting, feeding and grooming. The ferrets ranged in age from 8 months to 7.5 years of age over 21 females and 8 males. A home sampling kit was sent to the participants including material to safely collect and store ferret oral swabs. One participant had significant animal handling experience and performed all sample collection to standardize sampling procedures. Thirty oral swabs were collected and held in viral transport media in the participants’ freezer until the end of the study period. Frozen samples were directly transferred to a lab member and processed.

All samples were confirmed to have viable RNA by a preliminary screen for constitutively expressed ß-actin (Table 1). Each sample was then tested for evidence of active or recent SARS-CoV-2 infection with three established primer sets: ORF1b-nsp14 (20), Nucleocapsid (N) (14) and RNA-dependent RNA polymerase (RdRP) (21). All were below the limit of detection and determined to be negative for active or recent infection (Table 1).

**Table 1.** No evidence of SARS-CoV-2 infection in ferrets. Thirty samples from 29 ferret oral swabs were tested by semi-quantitative real time RT-PCR and ELISA. RT-PCR was performed on a StepOnePlus (ABI, Beverly, MA) with qScript XLT 1-Step RT-PCR ToughMix. Sample and RNA viability was confirmed by β-actin (ACTB). Three separate primers sets were used to test for SARS-CoV-2: ORF1b, N and RdRP; All SARS-CoV-2 results were under the limit of detection (LOD). Oral swabs were further tested for antibodies at a 1:5 dilution ELISA against recombinant protein A/G (Total IgG) or purified SARS-CoV-2 RBD (αRBD IgG). Plates were read on a BioTek Synergy 4 Multidetection plate reader (Winooski, VT). Positive cutoff was set at (μ + 3σ) of the negative controls (n=24).

| Ferret | ACTB | ORF1b | N | RdRP | Total IgG | αRBD IgG |
| --- | --- | --- | --- | --- | --- | --- |
| 1 | 33.036 | LOD | LOD | LOD | P | N |
| 2 | 28.120 | LOD | LOD | LOD | P | N |
| 3a | 27.954 | LOD | LOD | LOD | P | N |
| 3b | 28.945 | LOD | LOD | LOD | P | N |
| 4 | 26.230 | LOD | LOD | LOD | P | N |
| 5 | 29.067 | LOD | LOD | LOD | P | N |
| 6 | 29.729 | LOD | LOD | LOD | P | N |
| 7 | 29.360 | LOD | LOD | LOD | P | N |
| 8 | 26.755 | LOD | LOD | LOD | P | N |
| 9 | 33.049 | LOD | LOD | LOD | P | N |
| 10 | 32.820 | LOD | LOD | LOD | N | NA |
| 11 | 29.781 | LOD | LOD | LOD | P | N |
| 12 | 29.010 | LOD | LOD | LOD | P | N |
| 13 | 27.730 | LOD | LOD | LOD | N | NA |
| 14 | 32.163 | LOD | LOD | LOD | P | N |
| 15 | 30.230 | LOD | LOD | LOD | P | N |
| 16 | 27.861 | LOD | LOD | LOD | P | N |
| 17 | 27.701 | LOD | LOD | LOD | P | N |
| 18 | 27.687 | LOD | LOD | LOD | N | NA |
| 19 | 30.832 | LOD | LOD | LOD | N | NA |
| 20 | 31.758 | LOD | LOD | LOD | P | N |
| 21 | 31.758 | LOD | LOD | LOD | N | NA |
| 22 | 32.635 | LOD | LOD | LOD | P | N |
| 23 | 27.098 | LOD | LOD | LOD | P | N |
| 24 | 29.290 | LOD | LOD | LOD | P | N |
| 25 | 29.806 | LOD | LOD | LOD | N | NA |
| 26 | 35.042 | LOD | LOD | LOD | N | NA |
| 27 | 30.032 | LOD | LOD | LOD | P | N |
| 28 | 31.464 | LOD | LOD | LOD | P | N |
| 29 | 29.476 | LOD | LOD | LOD | P | N |

We further took advantage of salivary immunoglobulin, which has been shown to be highly sensitive and specific for SARS-CoV-2 testing (22). We tested samples for evidence of antibodies against SARS-CoV-2 surface glycoprotein receptor binding domain (RBD). Twenty-two ferrets (23 total samples) were confirmed to have measurable total IgG via binding to recombinant protein A/G but were all negative for binding to RBD (Table 2). Therefore, there is no evidence of viral infection or seroconversion in 29 ferrets living with two people with COVID-19.

### Identification of two mustelid-associated mutations in SARS-CoV-2 surface glycoprotein

Our observed household data support that there may be important barriers to natural infection in ferrets, however, ferrets have been shown to be susceptible to infection and onward transmission in experimental laboratory infections (9, 10, 18, 19). To further investigate this, we analyzed all currently available genomic sequences of SARS-CoV-2 viruses of naturally infected American minks and experimentally infected ferrets (32 sequences representing 24 animals, accessed: 2020-08-01). There are viral sequences available from two natural reverse zoonotic events in mink farms in Europe, which allowed us to infer founder-effect mutations versus acquired mutations of relevance to spillover (11). We identified three mutations of interest in the surface glycoprotein (S protein) coding sequence: N501T, D614G and S686G (Fig. 2A).

**Figure 2.**
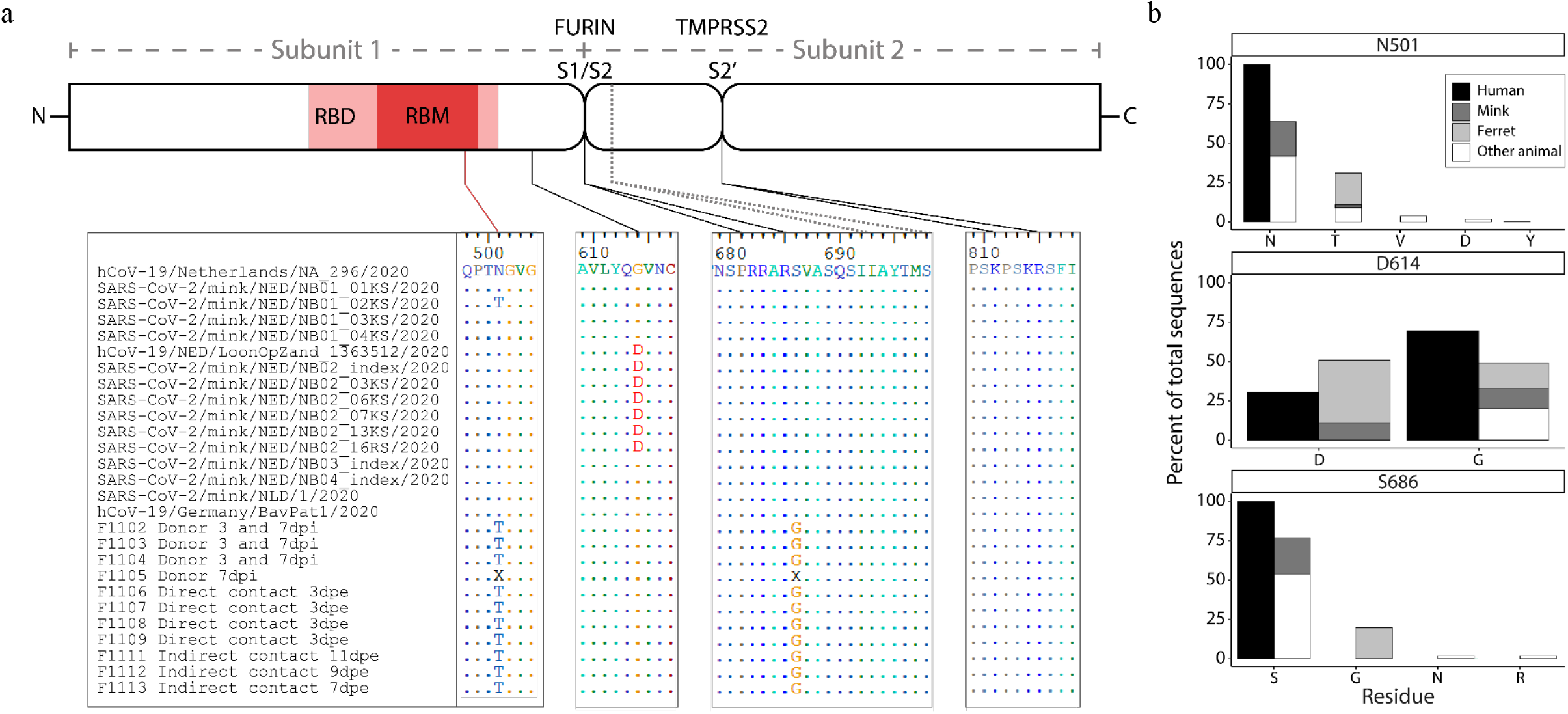
Mustelid-associated mutations in SARS-CoV-2 surface glycoprotein. All available SARS-CoV-2 surface glycoprotein (S) sequences from natural (mink) and experimental (ferret) infections were compared and three mutations identified. a) A schematic diagram (not to scale) of the S protein with Subunit 1, which is involved in host receptor protein attachment and Subunit 2, which is involved in host cell fusion. Mutation N501T is located in the receptor binding domain (RBD) and receptor binding motif (RBM), shown in red. Mutation D614G is located in Subunit 1 downstream of the RBD, and mutation S686G is located directly adjacent to the novel S1/S2 cleavage motif (PPAR↓**<u>S</u>**) processed by furin. A second S1/S2 cleavage site (IAY↓TMS) seen in SARS-CoV is conserved. The S2’ cleavage site (KPSKR↓S) processed by TMPRSS2 is also conserved. Viral amino acid sequences from regions of interest are shown below the schematic, and dots represent conserved residues using the top sequence as a reference (hCoV-19/Netherlands/NA_296/2020). Viruses from mink are separated into two clades from distinct farms (NB01 and NB02-4, respectively), and are preceded by the closest observed human sequence (hCoV-19/Netherlands) for reference. Experimentally infected ferrets are in the bottom half (F1102-1113). The sequence from the human inoculum (hCoV-19/Germany) is included for reference. Ferrets are separated into three groups: donors, which received direct inoculum; direct contact, which were housed with donors; and indirect contact, which were housed adjacent to donors without physical contact. Identical sequences were found from samples taken at 3 and 7 days post inoculation (dpi) in 3 of 4 donors. Donor F1105 exhibited two equivalent single nucleotide variants (A1502C and A2056G) resulting in N501/N501T and S686/S686G, respectively, and are not consensus-called (“X”) in those locations. b) 9,253 human-derived SARS-CoV-2 S protein sequences and 57 animal-derived SARS-CoV-2 or SARS-CoV-like virus S protein sequences were aligned to calculate percent amino acid representation at three positions: N501 (top), D614 (middle) and S686 (bottom).

First, N501T was observed in 11/11 experimentally infected ferrets (donor, direct and indirect contact), with an increasing proportion of the virome represented through the study period, supporting strong positive selection in ferrets (19). Only 1 of 13 mink viruses are N501T, which supports spontaneous mutation and natural selection in the population. The measured mutation rate calculated from the closest observed human-derived sequences in mink is very low, 4.2×10^−4^, so we asked if this specific mutation is otherwise common and not unique to mustelid infection. Of 9,049 high quality human-derived SARS-CoV-2 S genes, none exhibit the N501T mutation (Fig. 2B). However, N501T is seen in 5/17 pangolin-derived SARS-CoV-2-like viruses. Notably, the equivalent residue in SARS-CoV is a threonine (T487).

We observed a second conserved mutation, D614G, in one of the two mink clades and all ferrets. However, this mutation has become prevalent in the human population (D614, 30.5%; D614G, 69.5%, Fig. 2B) and was observed in the ferret human donor and mink farm’s closest observed ancestor (Fig. 2A). We conclude that D614G mutations are due to variation in the human population/donors and are not specifically associated with mustelid infection.

The third non-synonymous S protein mutation, S686G, was only observed in ferrets and is located at the P1’ serine residue directly adjacent to the novel S1/S2 polybasic cleavage site (PRRAR↓**<u>S</u>**) (Fig. 2A). This mutation is of special interest as this cleavage site partially distinguishes SARS-CoV-2 from other SARS-like viruses and allows immune evasion prior to receptor binding (23-25). Like N501T, S686G was observed in 11/11 ferrets and was a minority variant in the donor inoculum and increased proportional representation in the virome over time, suggesting positive selection (19). We found that no other human-derived viral sequence has been observed with this mutation (Fig. 2B). S686G has also not been observed in SARS-CoV-2-like viruses from other carnivores (naturally infected felines and canines), all of which retained the complete cleavage site and adjacent P1’ serine. All mustelid-derived viruses retained the second, downstream S1/S2 cleavage site motif (IAY↓TMS), as well as the S2’ TMPRSS2-processed cleavage site for fusion.

Host furin and furin-like proteases have been shown to cleave the S1/S2 polybasic cleavage site (3, 25, 26). P1’ residues are strongly favored to be serine in furin cleavage, and alternate residues are restricted by size and hydrophilicity due to their location in the furin binding pocket (27). Glycine is small but hydrophobic. We performed *in silico* analysis of the cleavage site to compare identical sequences that differed only at position 686 using PiTou 2.0 (28). PiTou scores are biologically meaningful prediction values of furin cleavage derived from binding strength and solvent accessibility and can be directly compared. S686 results in a PiTou score of 9.19633 while S686G results in a score of 6.92387. While both are predicted to be cleaved by furin, S686 is estimated to have stronger interactions in the binding pocket (P6-P2’). Therefore, S686G is an unfavorable substitution for furin cleavage.

We further performed phylogenetic analysis of the proprotein convertase family that cleave polybasic sites (PCSK1-7), including furin, and Cathepsin L in a number of mammals including *Mustela putorius furo* and the well-annotated *Mustela erminea*. However, we found no significant difference between ferrets, ermines and other carnivores.

## Discussion

Multiple studies have now demonstrated that ferrets may be directly infected by human-derived SARS-CoV-2 and, following infection, exhibit a 100% transmission rate via direct contact (9, 10, 18, 19). However, our data suggest that the initial barrier of human-to-ferret transmission may be higher than relevant for most household pets. We calculated that a sample size of 10 animals was sufficient to test the hypothesis that at least one ferret was infected, given an observed attack rate of 87% in mink farms (95% CI, 0.05) (29). In this natural experiment, all 29 ferrets had significant opportunities for direct contact with all other ferrets and had direct exposure to at least one, and likely two infectious people. While we were unable to collect human samples, current epidemiological knowledge of SARS-CoV-2 would lead to the conclusion that both adults had an infectious period with viral shedding (30, 31).

We found no evidence of SARS-CoV-2 transmission to ferrets based on RT-PCR and serology, a finding at odds with the high transmission rates observed in ferrets and mink and infectivity of SARS-CoV-2. Based on current knowledge of SARS-CoV-2 transmission and shedding in ferrets, we determined that our collection time points fell within the timeframe to obtain measurable viral RNA, even if transmission occurred on March 22, prior to any symptom onset in the household. However, it was important to perform additional antibody testing to address two concerns; first, that transmission could have occurred prior to March 22 and second, that the level of infection and viral shedding was so low as to be below collection and screening sensitivity. In either scenario, we still expected a robust antibody presence within days of initial infection but found no evidence of RBD-specific antibodies. Despite significant and prolonged exposure in the home, we have concluded that there is no evidence of SARS-CoV-2/COVID-19 human-to-ferret transmission in this household.

Notably, Ferret 12 (7yo) was euthanized on April 16, and had a history of adrenal disease, and Ferret 16 (7yo) died unexpectedly on April 20. Both were swabbed within four days of their deaths and we expect would have been RT-PCR or antibody positive had their deaths been related to SARS-CoV-2 infection.

Viral host receptors are often a key factor in determining host range. American minks and ferrets share 24 of 25 ACE2 residues with known viral S protein interactions, and we expect these species to have similar natural susceptibility (7). N501T is in the receptor binding motif of the SARS-CoV-2 surface glycoprotein, which interacts with ACE2 primarily at Y41, but also K353, G354 and D355 (32, 33). Of these, mustelids only differ from humans at ACE2 G354, and this site is also the only distinct residue between ferret (G354R) and American mink (G354H) (7). Mink have been naturally infected by virus without the N501T mutation and there have now been dozens of independent human-to-mink spillover events, therefore we do not expect that the ACE2 G354H mutation significantly limits infection. However, the appearance of N501T in all infected ferrets suggest ACE2 G354R may provide a host barrier to SARS-CoV-2 entry in ferrets. Additional work is needed to determine if N501T is a required adaptation for ferret transmission and, if so, if it affects transmission dynamics.

SARS-CoV-2 S protein S686G is another intriguing mutation as it lies directly adjacent to a motif that is likely to enhance virulence (25). To date, S686 is perfectly conserved in 9189/9189 human sequences, indicating strong purifying selection. S686G changes a neutral polar residue to a non-polar one, which we estimated to decrease furin efficiency. Furthermore, S686 completes a novel glycosaminoglycan (GAG)-binding motif (XBBXBX/PRRAR**<u>S</u>**) that enhances binding and the two flanking serines in the S1/S2 site (**<u>S</u>**PRRAR↓**<u>S</u>**V) have been shown to be permissive to host phosphorylation and consequent down regulation of furin activity, (26, 34). We were surprised to see evidence of positive selection over time for this potentially unfavorable mutation in ferrets as described by Richard *et al* for these reasons (19). If there is further evidence of S686G selection in experimentally or naturally infected ferrets, it is essential to fully investigate changes in viral fusion activity, kinetics and pathology to determine if ferrets are an appropriate model for human disease.

Our results suggest that virus and host genetic barriers significantly limit natural infection in ferrets, and these are only likely to be overcome by a concentrated and/or diverse inoculum of human-derived virus. To date, experimental ferret infections have been successful 6 × 10^5^ and 10^5.5^ TCID_50_, and at least one inoculum contained a minority of virus with the N501T and S686G variants (18, 19). These limitations and putative host-adaptations may negatively affect ferrets as a disease and/or transmission model and should be further investigated. We are, however, optimistic that the lack of spillover in this household supports that there is a very low risk of human-to-ferret SARS-CoV-2 transmission in domestic settings.

## Materials and methods

### Study enrollment and sample collection

The study participants were enrolled under a protocol approved by Tufts University Institutional and Animal Care and Use Committee and Health Sciences Institutional Review Board (#G2020-27). A self-administered sampling kit was sent to the enrollees’ residence with sterile standard polyester tipped applicators (Puritan, Guilford, ME), vials with 800ul M4RT viral transport media (Remel, Lenexa, KS), instructions, a data sheet and secondary containment bags. Oral swabs were obtained using gloves and a mask in the home and held in a home freezer until transfer to a lab member via a cooler.

### RNA extraction and RT-PCR

Samples were vortexed and 50ul aliquoted for MagPlate OMEGA extraction following manufacturer protocols. RNA was tested by semi-quantitative real time reverse transcription polymerase chain reaction (RT-PCR) on the StepOnePlus platform (ABI, Beverly, MA) with qScript XLT 1-Step RT-PCR ToughMix, using five primer sets: one for internal controls (ACTB) and three for SARS-CoV-2 (ORF1b, N1, E, RdRP). CoVERS-ACTB, F: GATGCAGAAGGAGATCAC, R: CTAGAAGCATTTGCGGTG, Probe: HEX-CTCCTGCTTGCTGATCCACA-TAM; HKU-ORF1, F: TGGGGYTTTACRGGTAACCT, R: AACRCGCTTAACAAAGCACTC, P: FAM-TAGTTGTGATGCWATCATGACTAG-TAM; 2019-nCoV_N1 [CDC], F: GACCCCAAAATCAGCGAAT, R: TCTGGTACTGCAGTTGAATCTG, P: FAM-ACCCCGCATTACGTTTGGTGGACC-TAM; RdRP_SARSr, F: GTGARATGGTCATGTGTGGCmGG, R: CARATGTTAAASACACTATTAGCAmTA, P: FAM-CAGGTGGAACCTCATCAGGAGATGC-TAM. All plates were run with negative VTM controls and positive control (NR-52285, Genomic RNA from SARS-Related Coronavirus 2, Isolate USA-WA1/2020, BEI Resources, Manassas, VA).

### ELISA

Oral swabs were tested for total IgG and IgG against SARS-CoV-2 receptor binding domain with minor modifications to an established protocol (35). Briefly, Immulon 2 HB plates were coated with 2μg/ml Pierce recombinant protein A/G (ThermoFisher catalog no: 77677) or purified SARS-CoV-2 receptor binding domain (provided by Florian Krammer, available as NR-52366, BEI Resources, Manassas, VA) and incubated 2 days at 4°C. After washing, plates were blocked with PBS supplemented with 0.1% Tween-20 (PBS-T) and 3% milk at room temperature for 2 hours. All samples were heat inactivated at 56°C for 1 hour. Ferret samples were diluted 1:5 in PBS-T with 1% milk. Positive controls were serum from S protein immunized alpacas (provided by Charles Shoemaker), and diluted 1:5 in PBS, then to final dilution of 1:50 in PBS-T with 1% milk. Following blocking, 100μl diluted samples were incubated at room temperature for 2 hours. Plates were washed and 50μl Pierce recombinant protein A/G with peroxidase (Thermo Fisher catalog no: 32490) added at 1:10,000 in PBS-T with 1% milk as a secondary and incubated 1 hour at room temperature. Plates were washed and developed for 10 minutes with SigmaFast OPD solution (Sigma-Aldrich catalog no: P9187), stopped with 50ul 3M HCl and read at an absorbance of 490nm on a BioTek Synergy 4 Multidetection plate reader (Winooski, VT). VTM was tested at 1:2 and 1:5 and confirmed to not affect results.

### Viral sequence collection and assembly

High quality SARS-CoV-2 surface glycoprotein sequences were curated using NCBI Virus and GISAID EpiCoV databases as follows. 9,664 full length S nucleotide sequences were collected from NCBI Virus and aligned using ClustalΩ 1.2.4. Sequences were trimmed to coding region sequence (CDS), translated and realigned. Sequences with >10% unknown residues were excluded. All non-human animal-derived SARS-CoV-2 and SARS-CoV-2-like viral sequences were collected from GISAID EpiCoV. To collect viral genomes from experimental ferret infection, sequencing reads were downloaded from 23 Illumina and Minion sequencing runs uploaded to NCBI Sequence Read Archive (PRJNA641813). Reads were confirmed to be post-quality control by Prinseq and mapped to the human donor sequence (hCoV-19/Germany/BavPat1/2020|EPI_ISL_406862|2020-01-28) using BWA (Illumina) and Pomoxis mini_align (Minion). Consensus was called using Samtools and replicate Illumina/Minion libraries were compared to confirm consistency.

### Mammalian gene collection, assembly and phylogenetic analysis

PCSK1-7 and CTSL sequences were collected from NCBI Orthologs from *Homo sapiens, Pan troglodytes, Sus scrofa, Ovis aries, Bos Taurus, Canis lupus familiaris, Vulpes vulpes, Felis catus, Panthera tigris altaica, Phoca vitulina, Mustela erminea, Myotis lucifugus, Eptesicus fuscus* and *Rousettus aegyptiacus. Mustela putorius furo* orthologs were inconsistent with related species by preliminary RAxML ortholog analysis. Seven publicly available RNAseq run from *Mustela putorius furo* (SRR11517721-SRR11517724, SRR391982, SRR391968, SRR391966) were downloaded and putative PCSK1-7/CTSL reads were extracted using BLAST. Reads were assembled using Pomoxis mini_assemble with ermine references. Reads were then mapped back to the proposed ferret assembly with BWA and well-supported consensus sequences were called using Samtools. Ortholog collections were analyzed using maximum-likelihood phylogenetics via RAxML (JTTγ using empirical base frequencies, 5000 bootstraps).

## Acknowledgements

This work was supported by NIAID grant HHSN272201400008C/AI/NIAID. We thank Dr. Florian Krammer for directly providing RBD and Dr. Charles Shoemaker for providing alpaca control serum for ELISAs. We thank Drs. Jennifer Graham and Elizabeth Rozanski for their assistance in connecting us to ferret-owners for the CoVERS study. We thank Lynne Christiansen and Mary Andersen for providing us masks to safely collect samples. We are especially grateful to our enrolled household who were exceptional participants during a challenging time.

